# Stored Elastic Bending Tension as a Mediator of Embryonic Body Folding

**DOI:** 10.1101/2024.04.02.587781

**Authors:** Mira Zaher, Ronit Yelin, Alaa A. Arraf, Julian Jadon, Manar Abboud Asleh, Sivan Goltzman, Lihi Shaulov, Dieter P. Reinhardt, Thomas M. Schultheiss

## Abstract

During development, amniote vertebrate embryos transform from a flat, multi-layered sheet into a three-dimensional cylindrical form, through ventral folding of the lateral sides of the sheet (the lateral plate, LP) and their fusion in the ventral midline. Although this basic aspect of vertebrate body plan formation has long been described, it is not understood at the mechanistic level. Each side of the LP is comprised of two tissue layers: a dorsal somatopleure (Sop) of pseudostratified coelomic epithelium, extracellular matrix (ECM) and ectoderm, and a ventral splanchnopleure (Spl) of similar construction except with endoderm instead of ectoderm. Using a chick embryo slice system we find that the flat stage is actually a poised balance of opposing elastic bending tensions, with dorsal bending tension in the Sop opposing ventral bending tension in the Spl. An intact extracellular matrix is required for generating the bending tensions, as localized enzymatic digestion of Sop or Spl ECM dissipates tension, while removal of the endodermal or ectodermal layers has no effect. As development proceeds, the Sop undergoes Epithelial-Mesenchymal Transition, ECM fragmentation and dissipation of dorsal bending tension, while the Spl ECM and ventral bending tension remain intact, thus changing the balance of bending forces in the LP to promote ventral folding. Consistent with these findings, interference with the elastic ECM component fibrillin in the Spl <u>in vivo</u> reduces stored bending tension and perturbs ventral body folding. A generalizable conceptual model is presented in which embryonic growth, in the context of specific embryonic geometrical constraints, leads to accumulation of bending tension in the LP ECM, which is harnessed to drive body folding.

## Introduction

A classic problem in embryonic morphogenesis is the generation of the cylindrical vertebrate body plan. The canonical vertebrate body has a “tube within a tube” structure, consisting of an internal gut tube surrounded by an external body wall, with multiple tissues and organs lying between these internal and external layers^1^. However, in many vertebrates, specifically in amniotes (a category that includes mammals, birds, and reptiles), the embryo starts out not as a cylinder but as a multi-layered flat sheet. The ectoderm, which will constitute the future external body wall, lies on the dorsal side of the sheet; the endoderm, which will form the gut tube, lies on the ventral side; and a mesodermal layer, which produces most of the internal body organs, lies between them. In order to achieve the final body shape, the lateral sides of the flat embryo (the lateral plates) bend ventrally, fusing in the ventral midline, to form the final tube-within-a-tube body structure (Fig. 1A-E)^2–4^. While a basic description of ventral body folding has been available for more than 100 years^3,5^, the mechanisms that underlie it are still not clear.

**Figure 1:**
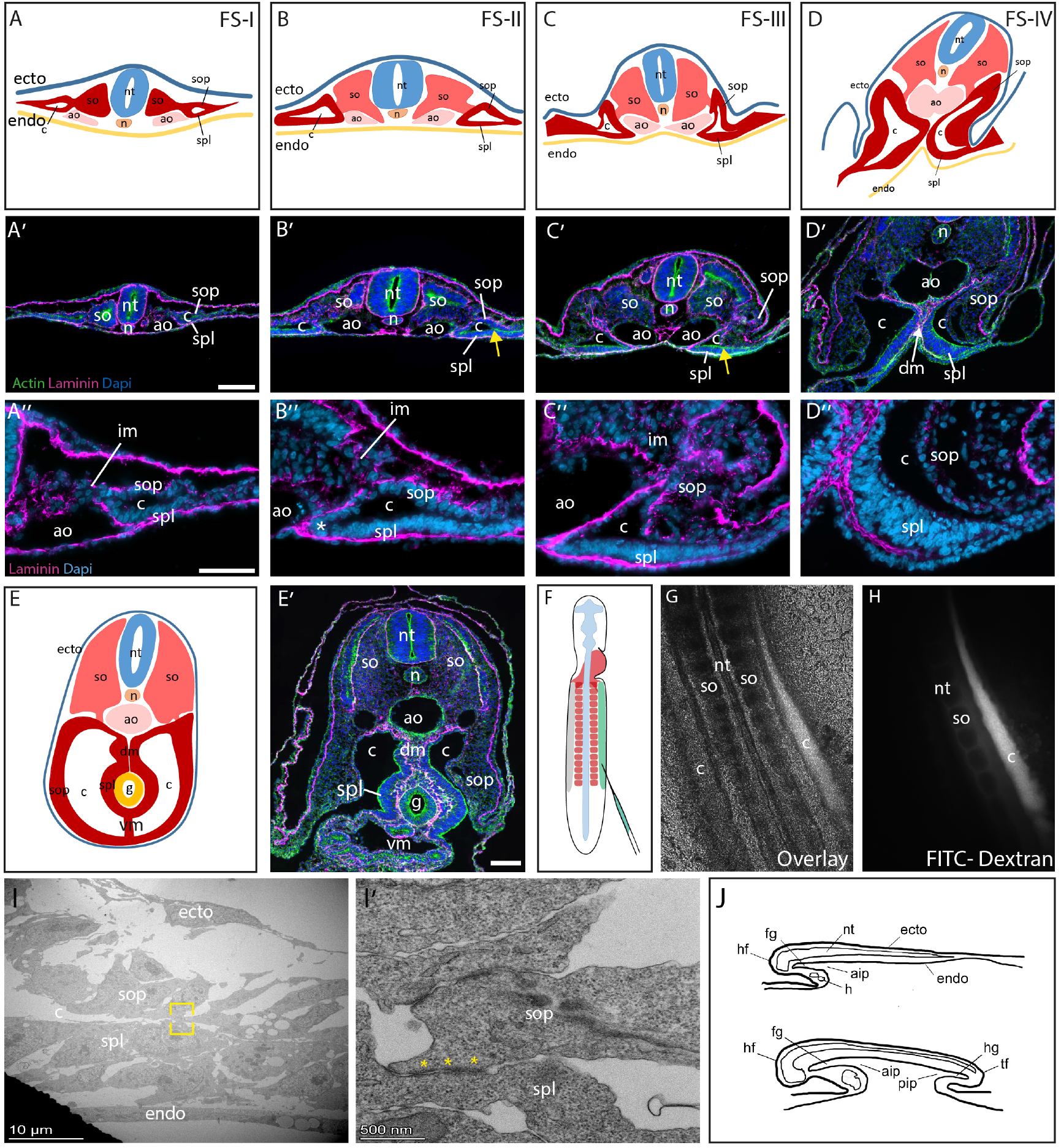
<u>Ventral Body Folding</u>. (A-E, A’-E’, A’’-D’’) Transformation from a flat disk to a cylinder through folding of the lateral plate, with closure of the gut tube and outer body wall. A’’-D’’ are enlargements of the right side of A’-D’. Arrows in B’, C’ mark lateral junction between Sop and Spl. Asterisk in B’’ marks Spl angle. (F-H) Injection of fluorescent Dextran into the coelomic cavity remains confined to the cavity. (I, I’) Transmission Electron Microscopy of the lateral junction between the Sop and Spl, showing junctional complexes at the joining point (asterisks). (J) Mid-sagittal diagram of embryo at HH Stage 10 (top) and 18 (bottom). Anterior is on left. Note progression of the AIP and formation of the PIP. AIP, anterior intestinal portal; Ao, aorta; c, coelom; dm, dorsal mesentery; ecto, ectoderm; endo, endoderm; fg, foregut; g, gut tube; h, heart; hg, nindgut; n, notochord; nt, neural tube; PIP, posterior intestinal portal; so, somite; sop, somatopleure; spl, splanchnopleure. Scale bars: A’, E’: 100μm; A’’: 50μm.

Body folding begins during neurula stages with formation of the head fold and foregut pocket (or anterior intestinal portal, AIP) at the anterior end of the flat embryo^4,6^ (Fig. 1J top). Somewhat later, the tail fold and hindgut pocket (posterior intestinal portal, PIP) form at the posterior end of the embryo^4,7^ (Fig. 1J bottom). Subsequently, as body folding progresses, the AIP and PIP advance posteriorly and anteriorly, respectively, until final closure is achieved in the midgut region with formation of the umbilicus^4,8,9^. Previous experimental studies on the initiation of folding in the AIP region of the chick embryo have suggested differing theories regarding the initiation of folding, including large scale traction forces from the retreating primitive streak^10^, and localized myosin-based contractility^6^, but the issue of mechanism is still unresolved. Studies on the posterior part of the chick embryo identified a role for a contractile force gradient under control of FGF signaling in extending the posterior gut pocket^7^, although the connection between these processes and general body folding is not yet clear.

Multiple^4,54,66,94,7,856^ mouse genetic mutants show defects in various stages of ventral body closure, including the transcription factors Tfap-2alpha^11^, Pitx2^12,13^, Gli3^14^, and Six4/5^15^, as well as the collagen protease BMP1^16^. BMP2 expression in the visceral/extraembryonic endoderm is required for initiation of ventral morphogenesis^17^, and Flk1 mutants, which lack all blood vessels, show defective body wall closure^18^. Defects in body folding have also been proposed to be at the root of human ventral body wall developmental anomalies including gastroschesis, omphalocoele, epispadias, and bladder exstrophy^19,20^. However, how these genetic mutations lead to problems with body wall closure, or how the human body wall closure defects are produced, is currently unknown.

Considerable progress has been made in recent years towards understanding mechanisms of morphogenesis^21–23^. Most of these studies have focused on cell-based sources of morphogenetic change, including force generation via myosin-based contractility^24,25^, regulation of cell replication^26–28^, or biomechanical properties of cells^29,30^. The Extracellular Matrix (ECM) has often been considered as a largely passive substrate for the morphologically active cells, although some studies have uncovered more active roles for the ECM during morphogenesis^31–35^.

The current study investigates body folding using the chick embryo model system. We describe the presence of stored elastic bending tension in the ECM of the lateral plate, and we present evidence that selective accumulation, storage and dissipation of this bending tension guides folding of both the gut tube and the external body wall. We suggest that regulation of stored ECM-dependent elastic tension within embryonic tissue layers may play a role in other large-scale morphogenetic movements.

## Results

### Tissue morphogenesis during ventral body folding

In the chick, as in other amniote vertebrates, body folding proceeds as a wave, beginning at the anterior end of the embryo with formation of the headfold at HH^36^ Stage 6 and proceeding posteriorly (Fig. 1A-E, J)^2^. At HH Stage 15, ventral folding is initiated at the posterior end of the embryo with formation of the tailfold, and folding proceeds in the anterior direction (Fig. 1J)^7^. The two waves of ventral folding eventually meet in the middle of the embryo to close the ventral body wall around the umbilicus. The current study focuses on folding in the mid-anterior embryonic region, between the heart and umbilicus.

In initial studies we documented the stages of body folding in sectioned embryos (Fig. 1A-E). In this figure, sections were taken at approximately the same axial level (somite 13) at successive stages of development. Because ventral folding occurs as a wave, each axial level passes through the stages of the folding process at a different chronological stage. Thus we will use the term “Folding Stage” or “FS” in order to describe the stages of ventral folding using a unified terminology independent of the particular axial level and chronological age.

At FS-I, the embryo is flat (Fig.1A-A’’). The lateral plate mesoderm consists of two pseudostratified epithelial layers: one adjacent to the ectoderm and the other adjacent to the endoderm. Classically, the ectoderm together with its adjacent mesoderm is called the somatopleure (Spl), while the endoderm together with its adjacent mesoderm is called the splanchnopleure (Sop). The apical sides of the splanchnopleuric and somatopleuric mesoderm face each other and are separated by the coelom, or embryonic body cavity, which at this stage is a thin fluid-filled space, and the pseudostratified epithelium that lines the coelomic space is called coelomic epithelium (CoEp). The medial boundary of the coelomic cavity is located where the lateral plate mesoderm meets the intermediate mesoderm (IM). The dorsal aorta are located at the junction of the somite, splanchnopleuric mesoderm, and endoderm.

By FS-II (Fig. 1B-B’’), several changes have occurred in the embryo as a whole and in the lateral plate in particular. The coelomic space has widened significantly due primarily to expansion of the medial wall of the coelom in the dorsal-ventral dimension. The two aortae have moved medially towards the midline and now lie ventral to the somites^2,37,38^. Together with the aortae, the ventral-medial border of the splanchnopleuric mesoderm also moves medially^2,37^, so that there is now an acute angle in the splanchnopleuric mesoderm at its leading, medial edge (Fig. 1B’’, asterisk). At this stage, the Spl is still relatively flat, while the Sop has adopted a domed shape indicative of the onset of folding in the Sop layer. The lateral extent of the coelom, where the Sop and Spl meet laterally (Fig. 1B’, arrow) marks the border between the embryonic and extraembryonic tissues. By FS-III (Fig. 1C-C’’), the Sop continues to bend and the Spl, while still largely flat, has thickened in the apical-basal dimension. The aortae and the ventral-medial border of the splanchnopleuric mesoderm have moved further medially, with the structures approaching their counterparts from the opposite side in the embryonic midline, ventral to the notochord, which has receded dorsally. By FS-IV (Fig. 1D-D’’), ventral folding of the Spl is under way. The aortae have met in the ventral midline and the splanchnopleuric mesoderm in the midline has lengthened to form the dorsal mesentery, which suspends the gut tube in the coelomic cavity^2,39^. Subsequently, the Spl from the two sides of the embryo fuse in the ventral midline to close the gut tube, followed subsequently by fusion of the Sop’s to close the external body wall (Fig. 1E,E’).

### The Somatopleure and Splanchnopleure are Mechanically Linked

In order to understand the mechanics of body folding, it was important to determine whether the Sop and Spl layers are mechanically linked. On the medial side of the coelomic cavity the two layers form a continuous epithelial layer where they join the intermediate mesoderm (Fig. 1A’’,B’’ asterisks). Thus it is clear that the two layers are linked on their medial sides. However, on the lateral side the situation was less certain. In particular, there are Sop and Spl layers in the extraembryonic tissues that are continuous with their intraembryonic counterparts (Fig. 1A’-C’), and there is an extraembryonic coelom between the two extraembryonic mesodermal layers. It was not clear to what extent the intraembryonic Sop and Spl layers interact with each other laterally in order to form an integrated mechanical unit.

We performed several studies to examine the lateral connection between the Sop and Spl layers at the lateral border of the coelomic cavity. First, fluorescently-labeled Dextran was injected into the embryonic coelomic cavity. Imaging of injected embryos revealed that the fluorescent dye remained in the embryonic coelomic cavity and did not leak into the extraembryonic space (Fig. 1F-H), indicting a tight seal at the lateral coelomic border. In addition, examination of the lateral coelomic border by Transmission Electron Microscopy (TEM) revealed the presence of multiple electron-dense junctional complexes between epithelial cells of the Sop and Spl (Fig. 1I-I’). These data indicate the existence of a strong connection between the Sop and Spl layers at the lateral intraembryonic coelomic border, and suggests that the two layers may be mechanically linked. Importantly, while in most regions of the lateral plate the apical surfaces of the Sop and Spl face each other across the coelomic cavity, at the lateral coelomic border the Sop and Spl are twisted with respect to each other, allowing for the formation of junctional complexes between the lateral edges of Sop and Spl cells (Fig. 1I’).

### Stored Elastic Tension in the Somatopleure and Splanchnopleure

In order to begin to explore the mechanical properties of the Sop and Spl, we developed an experimental slice system (Fig. 2). Full thickness slices approximately 200 microns in thickness were cut from living FS-I or II embryos and placed onto silicone-coated dishes in a cross-sectional orientation (Fig. 2A-C). Next, the lateral attachment between the Sop and Spl was severed mechanically using a microscalpel (Fig. 2D, E, G). It was found that immediately after performing this operation, the Sop recoiled strongly and quickly dorsally, while the Spl recoiled strongly and quickly ventrally (Fig. 2F, G, 3; Movie M1). Time-lapse imaging of the slices during the operation revealed that significant recoil of the Sop and Spl occurred within seconds after severing their lateral attachment (Fig. 3H, 32.0% +/- 10.8% of the equilibrium opening angle was attained within 1 second after cutting, n=8), with equilibrium positions attained after approximately 2 minutes (Fig. 3G). These data indicate that at FS-I and II, both the Sop and Spl layers are under significant bending tension and that these tensions are in opposing orientations. These finding also suggests that the relative flatness of the intact lateral plate mesoderm at these stages is not due to absence of bending tension, but to a balance of opposing tensions in the Sop and Spl combined with adhesion between the layers laterally.

**Figure 2:**
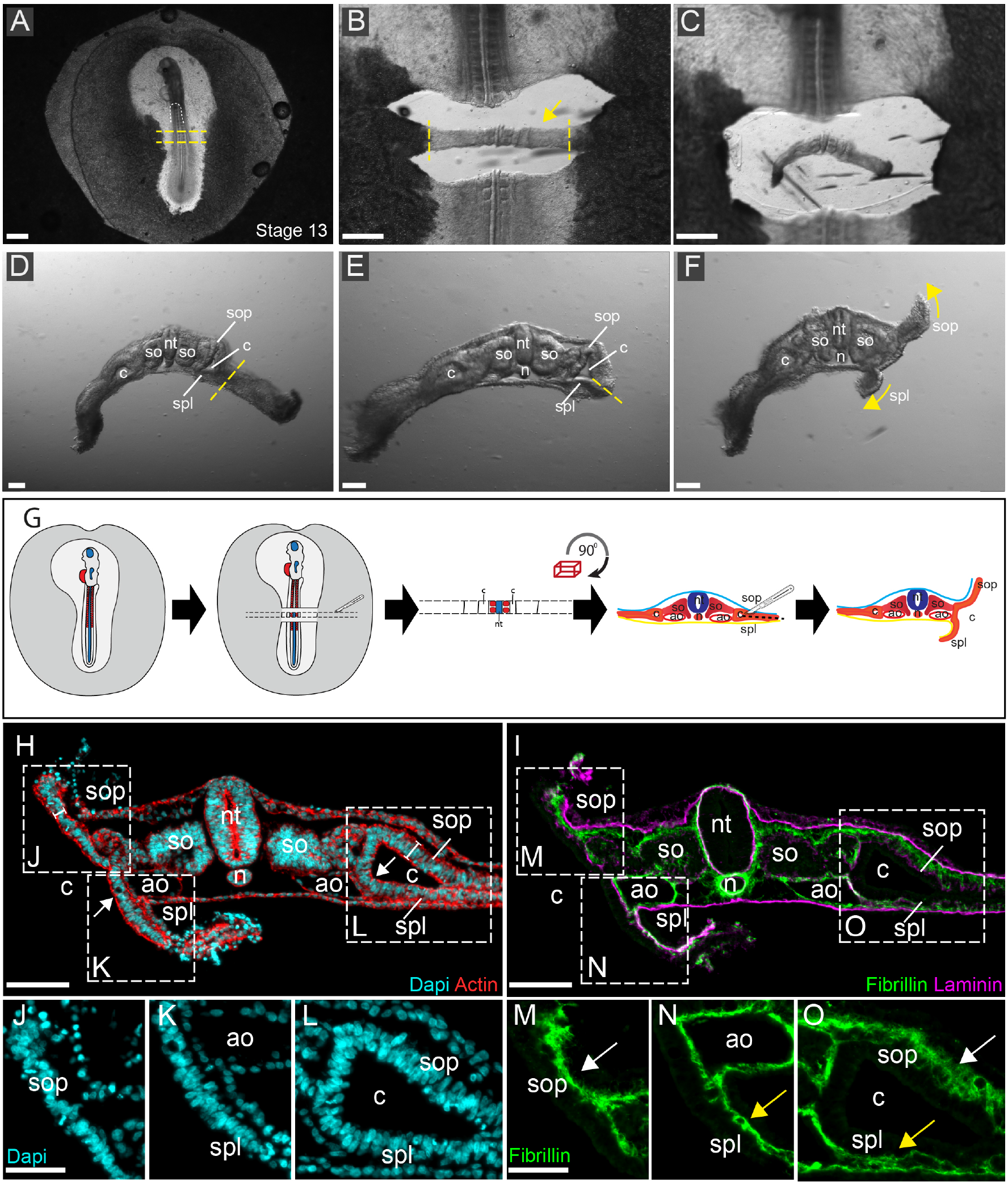
<u>Stored elastic bending tension in the Sop and Spl</u>. (A-G) Cutting of slice and removal from embryo (A-C), removal of extraembryonic tissue (D), cutting with knife to separate the Sop from the Spl (E), and rapid recoil of the Sop dorsally and Spl ventrally (F, yellow arrows) (n=68 embryos). Blue dashed lines indicate cutting sites. These steps are illustrated diagrammatically in G. (H-O) Histology of tissues after cutting. The connection between the Sop and Spl was cut on the left side. J, K, M, N are magnifications of the cut side, and L, O are are magnifications of the uncut side. Note straightening of the acute angles where the Sop and Spl join the medial coelomic wall (H, arrows), and thinning and loss of pseudostratification of the Sop and Spl on the cut side. The fibrillin on the cut side undergoes partial collapse (M-O, arrows). Abbreviations as in Fig. 1. Scale bars: A, B: 500μm; C-F, H, I: 100μm; J, M: 50μm.

**Figure 3:**
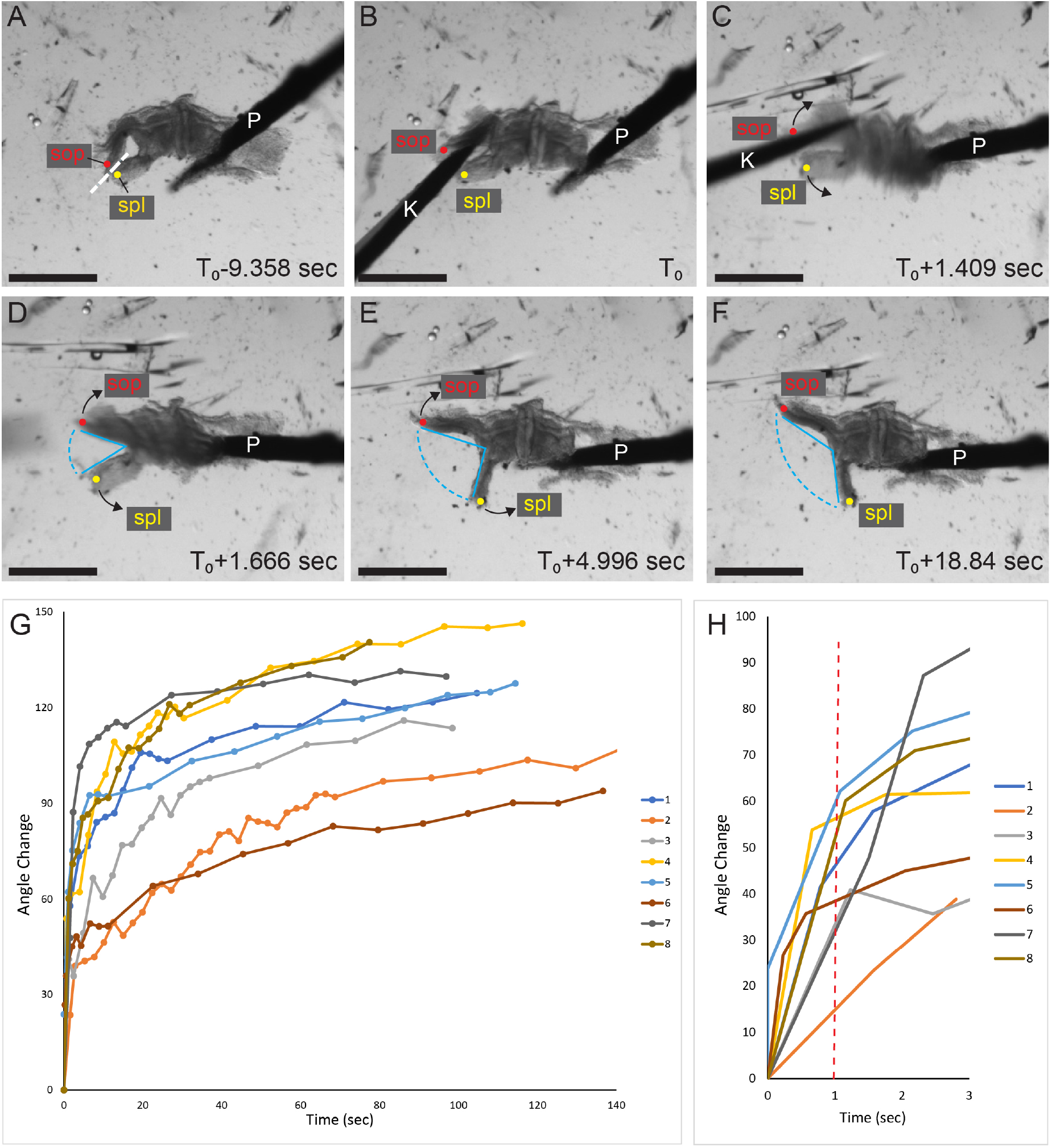
Elastic recoil after separating the Sop and Spl. (A-F) Time lapse movie taken while cutting the connection between the Sop and Spl. Blue bars indicate angle of opening between the Sop and Spl. (G, H) Graphs of opening of the angle between the Sop and Spl as a function of time (n=8). H is a magnification of the first 3 seconds after cutting. Line at 1 sec indicates timepoint for interpolated values, as described in Results and Methods. Scale Bars: 500μm.

Histological examination of slices after cutting revealed that the Sop and Spl have largely straightened, eliminating the acute bending angle that is seen on the control side (Fig. 2H arrows). This suggests that this acute angle may be the point in the intact slice where the Sop and Spl are at maximal bending strain and furthest from equilibrium. The apical actin foci present in the coelomic epithelium at this angle are maintained after cutting (Fig. 2H, arrows), suggesting that the opening of the angle is not due to loss of apical tension, and may instead be caused by changes in the basal zone. Release of the lateral connection between the Sop and Spl also resulted in thinning and loss of pseudostratification of the coelomic epithelium (compare Fig. 2J, K to Fig. 2L), indicating that the pseudostratified epithelium is normally under compressive strain, which is released upon severing of the mechanical linkage between the Sop and Spl layers. The ECM component Fibrillin-2 (Fbn2) transitions upon cutting from an extended meshwork to a more condensed, thinner arrangement (Fig. 2M-O), suggesting that the ECM is also under strain, and collapses upon relaxation of the applied stress that occurs after cutting of the lateral connection between the Sop and Spl.

### The ectoderm and endoderm do not contribute significantly to the elastic bending tension of the lateral plate

The Sop is comprised of several layers: the ectoderm, the coelomic epithelium, and the extracellular matrix and mesenchymal cells that lie between them (Fig. 1A-D). For the Spl the situation is similar, except that the external epithelial layer is the endoderm. In order to understand the nature of the stored elastic tension in the Sop and Spl, it was important to determine which component(s) of these layers store the tension. As a first step, we considered the roles of the ectoderm and endoderm. A patch ectoderm or endoderm was mechanically peeled from intact embryos using sharpened tungsten needles. Then, slices were cut through the peeled region and the lateral attachment between the Sop and Spl was severed as in Figure 2 (Fig. 4A-D). We observed that both the Sop and Spl recoiled normally despite lacking the ectodermal or endodermal layers (Fig. 4B, D). This indicates that elastic tension is not stored within these layers. Examination of histological sections of embryos in which the ectoderm had been removed (Fig. 4E-H) showed that the ectoderm had been cleanly removed (Fig. 4E- H asterisks), but that other components of the Sop remained intact, including the ECM as well as other cellular components of the layer (Fig. 4E-H’). Thus, the bending tension could reside in any of the components that remained after the removal of the ectoderm and endoderm.

**Figure 4:**
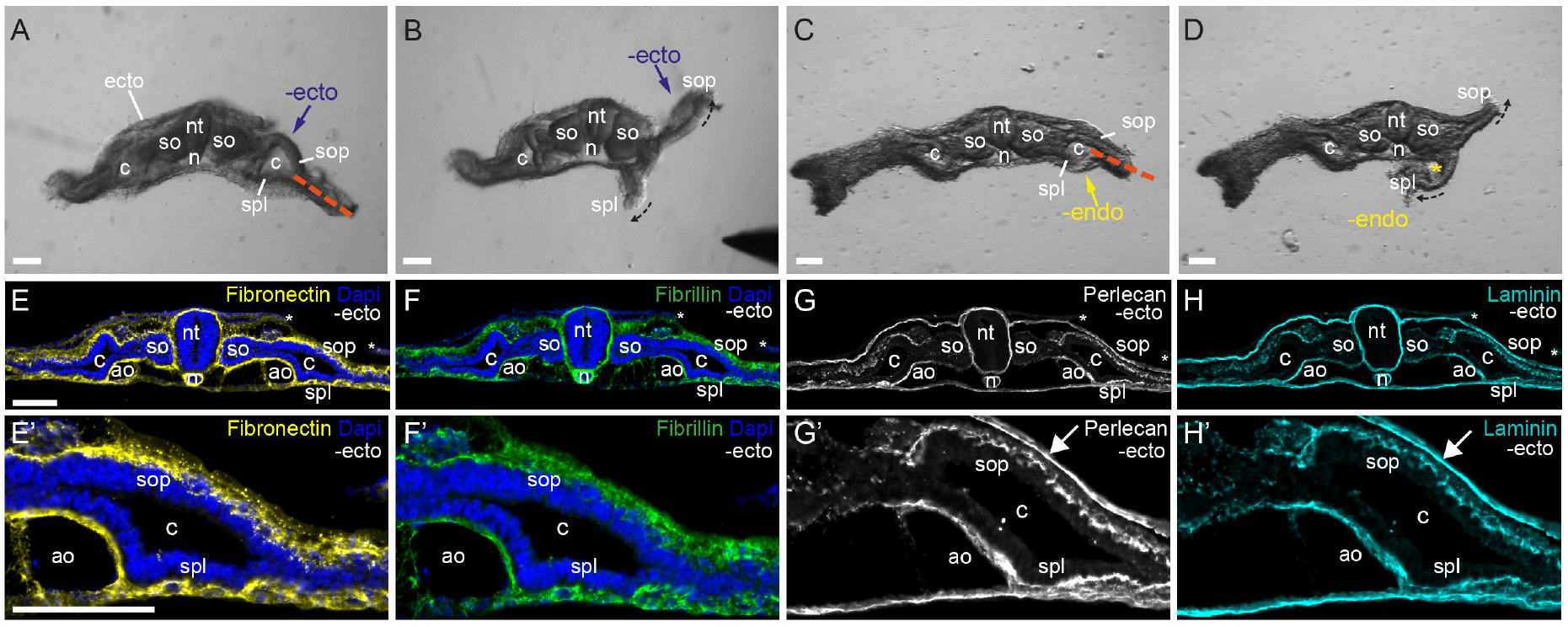
<u>Ectoderm and endoderm do not contribute to the elastic properties of the Sop and</u> <u>Spl</u>. (A-D) Removal of the Ectoderm (A, B) or endoderm (C, D) before cutting the connection between the Sop and Spl (red dashed line) did not affect the elastic recoil of these layers (ectoderm n=16, endoderm n=14). (E-H) Ectoderm was removed from a region on one side of the embryo. Edges of ectoderm are marked by asterisks. ECM of the Sop remains intact after ectoderm removal (compare left and right sides of the sections). Note that the basement membrane of the ectoderm also remains intact (G’, H’ arrows). Scale bars: 100μm.

### The extracellular matrix stores elastic bending tension

Next, we examined the role of the ECM in generating bending tension in the lateral plate (Fig. 5). First the ectoderm from FS-I or II embryos was peeled away in order to provide access to the subectodermal layers (Fig. 5A arrow; as described above, the ectoderm does not contribute to the elastic properties of the Sop). Then a solution of the protease Dispase was briefly applied to the exposed surface. The embryos were washed to remove the protease, slices were cut, and the lateral attachment between the Sop and Spl was severed. Under these conditions, there was a severe loss in recoil of the Sop when compared with embryos that had not been treated with Dispase (Fig. 5A, B), while recoil of the Spl was not affected. Similar results were seen when Dispase was applied to the Spl after removal of the endoderm (Fig. 5N- P’). This indicates that a Dispase-sensitive component of the Sop and Spl is required for storing bending tension.

**Figure 5:**
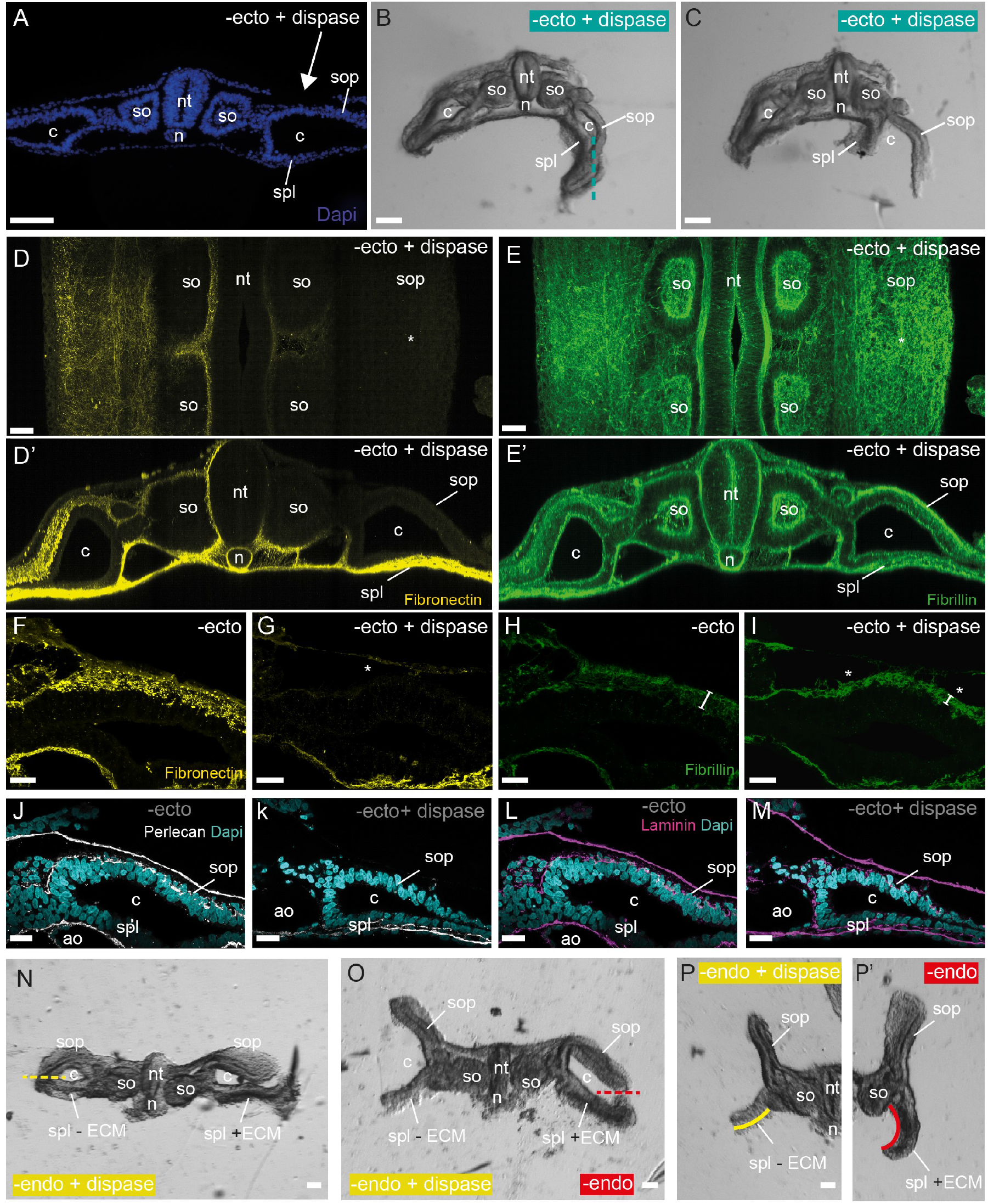
<u>Digestion of ECM dissipates stored elastic tension.</u> (A) Ectoderm was removed and a solution of Dispase was locally applied to the exposed Sop (arrow). (B-C) Slice before (B) after (C) cutting of the connection between the Sop and Spl (line in B indicates cut location), the Spl recoiled normally but the Sop did not recoil. (n=34) (D, E) Flat whole mount view (D,E) and reconstructed cross sections (D’, E’) showing digestion of FN (D, D’) and collapse and intensification of fibrillin (E, E’) on the Dispase-treated side. Histological sections of Dispase-treated (G, I, K, M) and control embryos (F, H, J, L) revealed loss of FN (F, G) and perlecan (J, K), and thinning (brackets in H) and condensation of Fbn2 (H, I) upon Dispase treatment, while laminin (L, M) was unaffected. (N-P) Endoderm was removed from the left side of the embryo and then the connection between the Sop and Spl was cut on both sides (dashed lines). The control side recoiled normally (P’). On the Dispase-treated side (P), the Sop recoiled strongly, while Spl recoil was impaired (compare yellow with red lines). (n=8) Scale bars: A-C, N-P: 100μm; D,E: 50μm; F-M: 20μm.

The Sop of Dispase-treated and control embryos was examined for expression and arrangement of ECM components. Fibronectin and the basement membrane associated protein Perlecan were locally decreased or absent in the regions treated with Dispase (Fig. 5, compare left and right sides of D, D’; also compare Fig. 5F, J with G, K). In contrast laminin was unaffected in the basement membranes of both the ectoderm (the ectoderm basement membrane was often not removed when the ectoderm was peeled away) as well as of the coelomic epithelium (Fig. 5L, M). The elastic extracellular matrix protein Fbn2 appeared to undergo a collapse, from an extended network in control embryos to a thinner, denser arrangement in Dispase-treated embryos (Fig. 5E, E’, H, I; note that H and I were photographed at identical exposure conditions). These results indicate that digestion of the ECM causes a loss of bending tension in the lateral plate, and thus suggest that the ECM is a major source of stored bending tension in the Sop and Spl layers.

### Changes in Elastic Tension in the Somatopleure and Splanchnopleure during the progression of body folding

Until now, the experiments were performed at FS-I or II, when folding of the lateral was in its early stages. In order to determine whether elastic tension in the Sop and Spl changes during the course of body wall folding, the slice experiment was repeated at different stages during the folding process. By FS-III, the Sop has adopted a domed shape, while the Spl is still straight (Fig. 6B). If the lateral attachment between the Sop and Spl is severed at this stage, the Spl still recoils ventrally as it did at FS-I or II (Fig. 6C). In contrast, the Sop remains static, does not recoil or straighten, and maintains its dome-shaped structure (Fig. 6C). This indicates that between Folding Stages II and III, the Sop loses its stored mechanical bending tension, while tension is maintained in the Spl. Thus, the balance of elastic bending tensions between the Sop and Spl changes between FS-II and FS-III, moving from a balanced relationship at FS-I and II to a ventrally-biased situation at FS-III.

**Figure 6:**
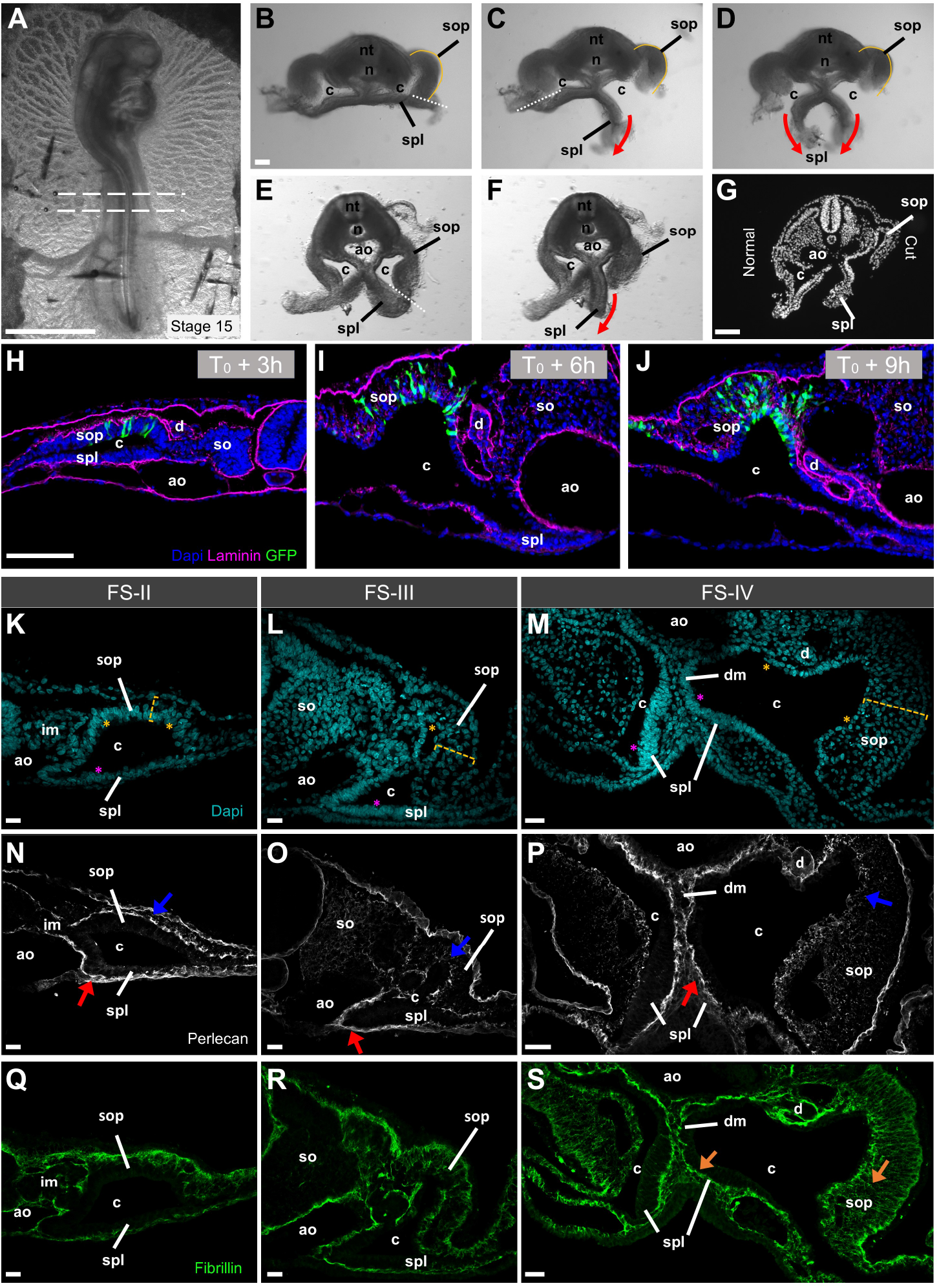
<u>Loss of Elastic Tension in the Sop at later stages and correlation with EMT.</u> Severing of the connection between the Sop and Spl at FS-II/III (B-D) or FS-IV (E-G) reveals that the Spl recoils ventrally, but the Sop does not recoil (n=30). A shows approximate level of sections of B-D). G is a section of at the same axial level as F stained with Dapi. After the right side was cut in C, the left side was cut (D). The right and left sides reacted similarly to cutting, indicating mechanical independence of the two sides. (H-J) The Sop undergoes EMT. The Sop was electroporated with a Gfp-expressing plasmid at FS-I and sections were taken at the indicated times. Note the movement of cells from the Sop epithelium into the newly forming mesenchyme. (K-S) Changes in the ECM during folding and EMT. Note the increase in thickness of the Sop (K-M yellow lines). The basement membrane associated molecule perlecan becomes fragmented in the Sop (N-P blue arrows) when compared to the Spl (red arrows). Fibrillin in the Sop becomes oriented perpendicular to the coelomic lining in the Sop, while it remains oriented parallel to the coelomic lining in the Spl (S, arrows). Scale Bars: A: 1.9mm; B, G-J: 100μm; K, L, N, O, Q, R: 20μm; M, S: 30μm; P: 40μm.

Similar results are obtained if the lateral attachment between the Sop and Spl is severed at FS-IV, when morphological bending is well underway in the Spl: the Spl recoils ventrally while the Sop maintains its morphology and does not recoil (Fig. 6F-G). This indicates that significant ventral elastic bending tension remains in the Spl throughout the stages when morphological folding is occurring.

As part of these experiments, the attachment between the Sop and the Spl at FS-III was severed on one side of the embryo and the Sop and the Spl layers were allowed to open until they reached equilibrium (Fig. 6C). Then, the attachment between the Sop and the Spl was severed on the other side of the same slice. In his case, the Spl on the second side recoiled with similar dynamics to the recoil that was observed on the first side (Fig. 6D). This indicates that the two lateral plates are mechanically isolated from each other, and that dissipation of tension on one side of the embryo does not affect the tension on the other side.

### Loss of elastic bending tension in the somatopleure correlates with Extracellular Matrix changes that occur during Epithelial-Mesenchyme Transition

In light of the change in elastic tension that occurs in the Sop between FS-II and III, we examined the tissue structure of the Sop more closely during these stages. Between FS-II and III, the somatopleuric coelomic epithelium begins to undergo Epithelial-Mesenchymal Transition (EMT), and cells leave the coelomic epithelium to generate the mesenchyme of the lateral body wall and the limbs^40^ (Fig. 6H-J). As EMT proceeds, the Sop basement membrane (containing perlecan and laminin) becomes fragmented as cells leave the confines of the CoEp to fill the space between the mesoderm and ectodermal layers (Fig. 6 N-P and Sup. Fig. S1), and fibronectin fibrils also become fragmented (Sup. Fig. S1). The fibrillin fibrils in the Sop become reoriented such that instead of being aligned parallel to the coelomic lining they become aligned perpendicularly to it, on an axis that stretches from the coelom to the ectoderm (Fig. 6Q-S). The Sop mesoderm layer becomes thicker and the cell density is reduced (Fig. 6K- M). The changes in Sop ECM pattern, combined with the finding that enzymatic disruption of the matrix causes a loss in elastic tension in the layer (Fig. 5B), suggests that the changes in the ECM that occur as a result of EMT may drive the changes in elastic properties of the Sop between FS-II and III. In contrast, the Spl CoEp remains largely intact during these stages (Fig. 6 G, K-S, Sup. Fig. S1), with intact basement membrane and fibrillar components and maintenance of fibrillin fiber orientation parallel to the coelomic lining, correlating with the maintained elastic properties of the Spl. In addition, cellular density of the splanchnopleuric CoEp layer increases (Fig. 6 G; asterisks in L, M), indicating maintenance of compressive forces in the Spl.

### Intact Extracellular Matrix is required for body folding in vivo

Finally, we investigated whether disturbance of the lateral plate ECM would influence body folding in vivo. We chose to begin by interfering with the function of the ECM component Fbn2^41^. Fibrillin microfibrils are an important component of the elastic properties of tissues and serve as templates for subsequent deposition of elastin^42^. A C-terminal truncated form of human Fbn2 (dnFbn2), which has been reported to interfere with fibrillin fibril formation^43^ (Fig. 7E), was electroporated into the Spl of embryos at FS-I/II (human chick Fbn2 are 95% homologous in this C-terminal fragment). Examination of embryos 24 hours after electroporation stained with an antibody that recognizes chick but not human Fbn2 (Sup. Fig. S2) shows diminution of Fbn2 immunostaining in electroporated areas, indicating that the Fbn2 construct is interfering with the fibrillin network (Fig. 7G, H, compare to control in I). We compared the morphology of dnFbn2-electroporated with control-electroporated embryos. Experimental and control embryos were always electroporated on the same side, in order to control for natural asymmetries that exist between the left and right sides of the lateral plate at these stages. We observed a lengthening of the Spl in dnFbn2-electroporated embryos (Fig. 7A-D, F), suggesting a disturbance in the ability of the Spl to resist stretching forces. In addition, we frequently observed that the Sop exhibited less extensive folding (Fig. 7A-D, arrows). Using the neural tube as an internal landmark, it can be seen that the dorsal-most point in the Sop has not moved as far ventrally in dnFbn2 as compared to control electroporated embryos (Fig. 7A-D, yellow lines). Since the dnFbn2 construct was electroporated into the Spl, the changes in the Sop must be secondary effects, likely via the mechanical linkage between the Sop and Spl (Fig. 1F-I), with elongation of the Spl leading to loss of medial pulling force on the Sop and defects in ventral Sop bending. These data support the proposition that an intact ECM network in the Spl is required for proper body wall folding in vivo.

**Figure 7:**
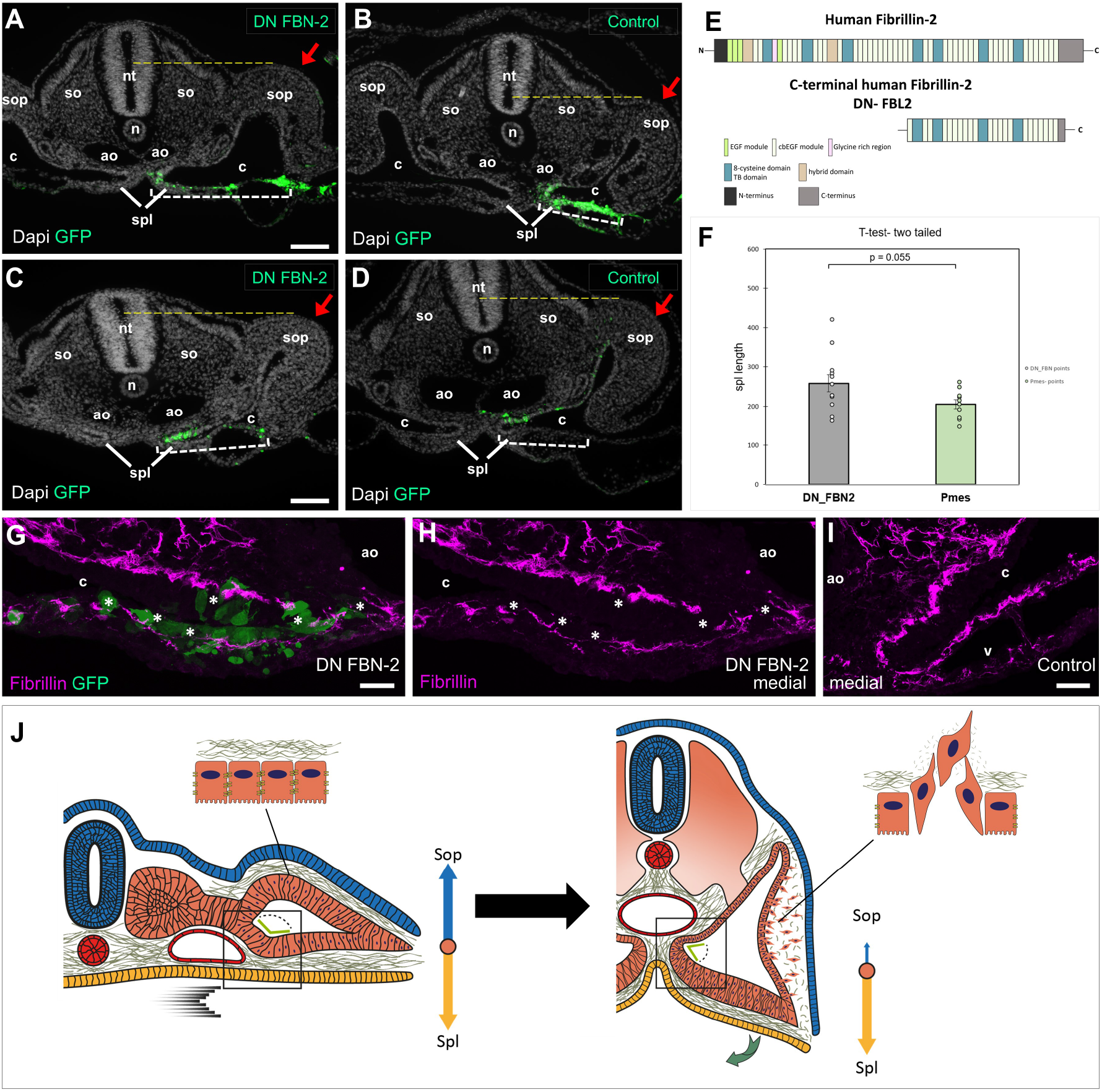
<u>Fibrillin disruption disturbs folding morphogenesis in vivo</u>. Truncated Fbn2 (dnFbn2; A, C, E) or control Gfp-expressing plasmid (B, D) was electroporated in the Spl. After 24 hours, the Spl was longer in dnFbn2 treated vs. control embryos (brackets in A-D; statistics in F, mean Spl length upon dnFbn2 electroporation 258 +/- 22μm [SEM], n=12 embryos vs. 205+/- 12μm, n=10 in control; p=0.55). Morphology of the Sop was often abnormal in dnFbn2 embryos, with the Sop exhibiting less ventral folding when compared to controls (arrows). Yellow lines show the relationship of the most dorsal point of the Sop to the neural tube, indicating less ventral Sop folding upon dnFbn2 embryos. (G-I) Electroporation of dnFbn2 caused fragmentation and lower staining levels of Fbn2 (G, H) when compared to the unelectroporated side of the same section (I). Images were captured under identical conditions. Scale Bars: A, C: 100μm; G: 20μm; I: 15μm. (J) Diagram of transition from a flat to a bending state, showing EMT and fragmentation of the ECM in the Sop, change in the angle (green bars) and medial movement (black arrows) of the Spl, and changes in the relative strength of bending tensions in the Sop and Spl (blue and yellow arrows). Boxes indicate equivalent areas in flat (left) and folding (right) stages. See text for details.

## Discussion

The current study revealed a system of stored ECM-based elastic bending tension in the embryonic lateral plate that can be harnessed for large scale morphological movements such as body folding.

In order to build up elastic tension, an elastic material must experience strain from an applied stress. Examination of the tissue arrangements in the lateral plate in the flat embryo stage reveals the existence of bends in the Sop and Spl where they take part in formation of the medial coelomic wall (Fig. 7 J, green bars). These bends are under tension, as is revealed when the lateral attachments between the Sop and Spl are severed; the Sop and Spl are now free to move, and they quickly recoil to eliminate the bend and form a straight line (Fig. 2, 3). We hypothesize that these bends occur, and tension is accumulated, through natural growth of the embryo. The Spl is attached to the endoderm through most of its length. It is also connected to the intermediate mesoderm and Sop at its medial and lateral ends, respectively, and to the aorta in middle (Fig. 7J; boxed area). As development proceeds, growth of the central part of the embryo in the dorsal-ventral dimension, including the somites, causes a lengthening of the medial coelomic wall. Because of the adhesion between the Spl and the aorta, and of both tissues with the endoderm, this causes bending strain in the Spl in the region of the aorta. Medial migration of the aorta and attached Spl to form the dorsal mesentery^2^ increases the acuteness of the bending angle in the Spl and likely augments Spl bending strain (Fig. 1A-C’, 7J, green bars). A similar process occurs in the Sop, with the nephric duct and ectoderm in place of the aorta and endoderm, respectively. In the Spl, this strain creates elastic bending tension that can be harnessed for ventral folding of the lateral plate, while in the Sop it is dissipated through changes in the ECM (Fig. 7; see also below). Thus, natural growth of the embryo, together with basic features of the embryonic anatomy, such as adhesion between key tissues, generates tension that can drive subsequent stages of embryonic morphogenesis. Accumulation of tension through differential tissue growth has been invoked as a source of energy to perform morphogenetic work in other contexts, for example to explain gut looping as a result of stored tension in the dorsal mesentery^39^.

The current study found that the flat lateral plate (FS-I) is not a neutral state lacking bending tension, but rather a dynamic balance in which the ventrally oriented Spl bending tension is opposed by a dorsally oriented Sop bending tension. In order to generate morphological tissue movement, this balance must be disrupted. This appears to occur, at least in part, through EMT-associated changes occurring in the Sop beginning between FS-II and III, which fragment the Sop basement membrane and ECM network. These changes are likely to be at least partly responsible for the reduction in Sop elastic bending tension observed at these stages. As a result of the change in the balance between the Sop and Spl stored tensions, the equation becomes tilted in favor of ventral bending (Fig. 7J). Thus, EMT not only produces local changes in the properties of the Sop to generate the Sop-derived somatic and limb bud mesenchyme^40^, but also facilitates ventral bending of the Spl because of the mechanical linkage between the two layers.

In the search for mechanisms that drive morphogenetic change, most studies have focused on cellular mechanisms, including contractility, growth, and physical cellular properties such as adhesion, stiffness, and fluidity^24–26,29^. The ECM has been much less studied as an active participant in shaping morphogenesis, although some recent studies have uncovered more active roles for ECM^31–35^. The current study found that an intact ECM is required for maintenance of tension within the Sop and Spl, and that interference with the ECM component fibrillin produces defects in the early stages of body folding in vivo. Elasticity of the ECM is well known for its role in short term storage and release of tension, as in the vessel wall of large arteries that undergo pulsatile flow^44^. In contrast, a role for ECM in long term storage of elastic energy that is harnessed for morphogenesis, as in the current case, is not well documented. While fibrillin, because of its elastic properties^45^, is a good candidate for mediating the mechanical properties of the lateral plate, ^44^other ECM components may also contribute to the stored tension. Epithelial basement membranes (BMs) have elastic properties^46^ and BM strain may contribute to stored bending tension in the lateral plate. Molecules such as perlecan, which was digested by Dispase (Fig. 5J, K) and which links the BM to the fibrillar ECM network^47^, may also play a role. As the ECM is an interconnected network^48^, it is likely that multiple components contribute to its elastic bending properties.

While the current study identifies stored ECM elastic tension as a contributor to the forces that generate body folding, other processes are likely to be involved. In particular, tension generated by actin-myosin contraction may also play a role in driving ventral bending. Cell-generated tension has been found to drive morphogenetic events in many other systems^7,24,49–52^. Bending CoEp cells exhibit marked lengthening in the apical-basal axis and shortening in the lateral axes (Fig. 1D’) that could be indicative of tightening of circumferential actomyosin belts that may aid in straightening the acute splanchnopleuric angle (Fig. 1C’, D’, 7J) and promote ventral bending^6,23,48–51^. Indeed, there are likely to be important connections between contractility and the ECM, since the cytoskeleton is connected to the ECM via integrin receptors and contractility and ECM assembly have important mutual influences on each other^53,54^. The relationship between ECM tension and CoEp contractility during folding will be an important area for future studies.

It is important to keep in mind that ventral folding occurs as a wave that moves from anterior to posterior in the anterior part of the embryo (Fig. 1J)^10^ (and from posterior to anterior in the posterior embryo^7^). Regions that are at more advanced stages in the folding process may aid the folding of regions that are less far along. This notion is supported by classic experiments which found that a midline incision in the AIP completely stops the folding wave^55,56^. Thus, the force vectors (and corresponding ECM strain vectors) that drive ventral folding likely do not lie solely along the dorsal-ventral axis but also incorporate an anterior-posterior dimension. In the future it will be important to investigate the cellular processes occurring at the ventral midline that integrate the folding tissues from the two sides of the embryo, and to understand how these processes facilitate the continuation of the folding wave.

Marfan syndrome, which is caused by mutations in fibrillin-1^57^, has ventral body wall phenotypes^58^. In addition, several mouse mutants exhibit strong ventral wall body closure defects^11–16,18^. The human congenital defects omphalocele, gastroschesis, epispadias, and bladder exstrophy exhibit faulty closure of the ventral body wall^19,20^. In all these cases, the mechanisms for producing ventral wall defects are not known. The current study suggests that it could be useful to investigate a potential role for stored ECM-dependent elastic bending tension in producing the ventral body wall defects seen in these scenarios. Finally, many other large-scale tissue bending and folding events occur during development whose mechanisms are not well understood, including formation of the broad ligament and fusion of the Mullerian tubes to form the uterus^59^; formation of the diaphragm from tissue folds in the coelomic wall^60,61^; and return of intestinal folds to the body cavity during gut morphogenesis^62^ to name just a few. The current study, together with past studies of morphogenetic process such as gut looping^34,39,63^, suggest that consideration of stored elastic bending tension, and the role of the ECM in its production and regulation, may provide new insights into a broad range of large-scale morphogenetic processes, and into developmental anomalies that arise from defects in these processes.

## Acknowledgements

This work was supported by grants to T.M.S from the Israel Science Foundation (1463/16, 1528/22), the Israel Cancer Research Fund, and the Rappaport Family Foundation.

## Materials and Methods

### Chicken embryos

Studies were carried out on fertilized Hyline strain chicken eggs (Moshav Orot, Israel) and incubated at 38.5 degrees C in a humidified incubator until the desired stage.

### Expression plasmids

pMES drives constitutive expression of an inserted gene and co-expresses eGFP under an IRES element^64^. pMES-2 was constructed by addition of a group of unique <u>enzyme</u> sites (NotI, RsrII, AgeI, <u>EcoRV</u>, Eco53kI, SacI, <u>XhoI</u>, AflII, MluI) into the polylinker site into the original pMES plasmid. DN-Fbn2 was constructed subcloning the sequence coding for the C-terminal half of human fibrillin-2 (Asp1531–Lys2771) with a C-terminus 6xHis tag into pMES-2.

### Immunofluorescence

Embryos were fixed in 4% Paraformaldehyde in PBS for 1 hour at Room Temperature or overnight at 4 degrees, embedded in 15% Sucrose/7.5% Gelatin/PBS, and cryosectioned at 10μm and 20μm thickness. Sections were permeabilized with 0.25% Triton X-100 in PBS and incubated overnight at 4 degrees C with primary antibodies in blocking solution (1% Bovine Serum Albumin, 2% Goat Serum, 0.2% Tween-20 in PBS). The following primary antibodies were used: mouse anti-GFP (1:500, Molecular Probes 3E6), rabbit anti-GFP (1:1000, Molecular Probes a-6455), chicken anti-GFP (1:500, Abcam 13970), mouse anti-Laminin (1:10, Developmental Studies Hybridoma Bank clone 3H11), rabbit anti-Laminin (1:300, Sigma-Aldrich L9393), mouse anti-Perlecan (1:10 Developmental Studies Hybridoma Bank clone 5C9), mouse anti Fibrillin-2 (1:10, Developmental Studies Hybridoma Bank clone JB3), rabbit anti-Fibronectin (1:400, Sigma-Aldrich F3648). Actin filaments were labeled with ActinGreen488 (1:25, Invitrogen R37110). Sections were then washed 3 times for 5 minutes each in 1XPBS and then incubated with secondary antibody for 2 hours at Room Temperature in a humidified chamber. Secondary antibodies used were Alexa488, Cy3, or Alex647-labeled Fab fragments (1:250, Jackson ImmunoResearch). Sections were washed 3 times for 5 minutes each in 1XPBS. Dapi (Sigma) was added at 1μg/ml during the second wash to mark nuclei. Stained sections were mounted using Fluorescence Mounting Medium (DAKO S302380) and Imaged with either a Zeiss Axioimager M1 widefield epifluorescence microscope and a Qimaging ExiBlue monochrome camera, or a Zeiss LSM880 confocal microscope. Image overlays were performed with ImagePro Plus software (Mediacy) or Imaris (Oxford Instruments) software.

### FITC-Dextran

Fluorescein isothiocyanate – dextran (FITC-Dextran; Sigma) was injected directly into the coelomic cavity in HH St. 12-13 embryos using a mouth pipette and pulled glass <u>microelectrode</u>. Embryos were imaged using a Zeiss Axioimager M1 widefield epifluorescence microscope and a Qimaging ExiBlue monochrome camera.

### Transmission Electron Microscopy (TEM) Imaging

HH Stage 12-18 Embryos were collected on Whatman paper rings and fixed in 3mm plates with 2% glutaraldehyde and 2% paraformaldehyde in 0.1 M Sodium Cacodylate buffer pH 7.4 containing 5 mM CaCl2 for 2 hours at Room Temperature (RT), then cleaned and moved to a glass vial with fresh fixation solution for 30 minutes at RT. Following fixation, embryos were washed with Sodium Cacodylate buffer, post-fixed and stained for 1 hour on ice with 1% Osmium TetraOxide, 0.5% Potassium hexacyanoferrate, 0.5% Potassium dichromate in 0.1M Cacodylate buffer containing 2 mM CaCl2, following which the samples were en-block stained with 1% uranyl acetate for 1 hour on ice. Then the tissue was dehydrated in a graded ethanol series and embedded in Epon812 (Electron Microscopy Sciences). 75 nm ultrathin sections were cut with a ultramicrotome UC7 (Leica), transferred to copper grids and viewed using a Talos L120C Transmission Electron Microscope at accelerating voltage of 120 keV.

### In vitro slice system

Chicken eggs were incubated at 38 degrees C until HH St. 12-17. Embryos were collected on Whatman paper rings and placed dorsal side up in a silicon dish (Sylgard) with 1ml of 1XPBS. Whole embryo slices approximately 200-300 microns thick (1.5-2 somites thick) are cut by hand using a microscalpel (Feather). Two cuts were made horizontally followed by two vertically in order to free the slice from the rest of the embryo. The free slice was then rotated in 90° and placed on the Sylcard dish in a cross-sectional orientation. In some cases, the slice was secured with insect pins (Fine Science Tools; see Fig. 3). Subsequently, a knife was used to make cuts in selected layers and the morphological changes in the slice were recorded by transmitted illumination using a Leica M205 stereomicroscope with DFC360FX camera and Leica LAS software. Peeling of tissues (ectoderm or endoderm) was performed mechanically with sharpened tungsten needles. After manipulation, some slices were fixed in 4% Paraformaldehyde/PBS and processed for immunofluorescence.

### Dispase

After removing the ectoderm or endoderm mechanically, 10µl of ECM degrading enzyme Dispase-ii (Roche) was applied to the exposed mesodermal surface for 5 minutes, washed three times using 0.5ml 1XPBS, before preparing the 200mn slices. In parallel, embryo slices were fixed and examined by immunofluorescence in order to assess the degree of digestion of the ECM by the enzyme treatments.

### In ovo electroporation

For coelom electroporations, chicken eggs were incubated at 38 degrees C. until HH St. 12-13. Embryos were staged according to Hamburger and Hamilton^65^. A circular window was made in the eggshell, and an Indian ink (Pelikan)/PBS solution (1:10) was injected into the subgerminal cavity to visualize the embryo. A solution of plasmid <u>DNA</u> and Fast green (0.1%) was injected into the coelomic cavity at the level of the last somite, or into the somitocoele of the 3 most posterior somites, using a Nanoinjector III (Drummond) and pulled glass microelectrode. Single plasmids were injected at the following concentrations: pMES-2 = 2400 ng; DN-FBN2 = 2400 ng. Two tungsten wire electrodes were placed between the vitelline membrane and the ectoderm, and embryos were pulsed 2-3 times for 25 msec at 12V, 1 sec interval using a BTX 830 electroporator. Embryos were sealed with beeswax and reincubated overnight until HH St. 16–17 and fixed in 4% Paraformaldehyde in PBS.

### Quantification and statistical analysis

*Opening angle:* The opening angle of eight different embryos in different time points was measured in ImageJ^66^. The vertex of the angle was positioned at the medial angle of the spl coelomic epithelium, with one arm ending along the endoderm and ending below the center of the notochord, and the other at the cut end of the spl coelomic epithelium. The angle change was calculated by subtracting each angle from the angle at T0, each embryo was plotted in a graph using Excel as shown in Figure 3. To calculate the percentage of the angle change, the angle change at each time point was divided by the angle at the final timepoint. Embryo data points were inserted into Matlab, the data was fitted to a 2^nd^ degree exponential function using a Curve Fitting plugin, and the value for X=1 sec (X- time in sec, Y- percentage of angle change) was extracted. The interpolated values for X=1 were used to calculate the mean and the standard deviation.

*Spl coelomic epithelium length:* To measure the length of the Spl one end was fixed at the medial angle of the Spl coelomic epithelium and the other at the lateral connection between the spl and sop coelomic epithelium. Three measurements were taken for each embryo and the mean of the three measurements was calculated. Two-Sample T-Test was used to compare the means, and STD of the Error was calculated.

---

**Supplemental Figure S1:**
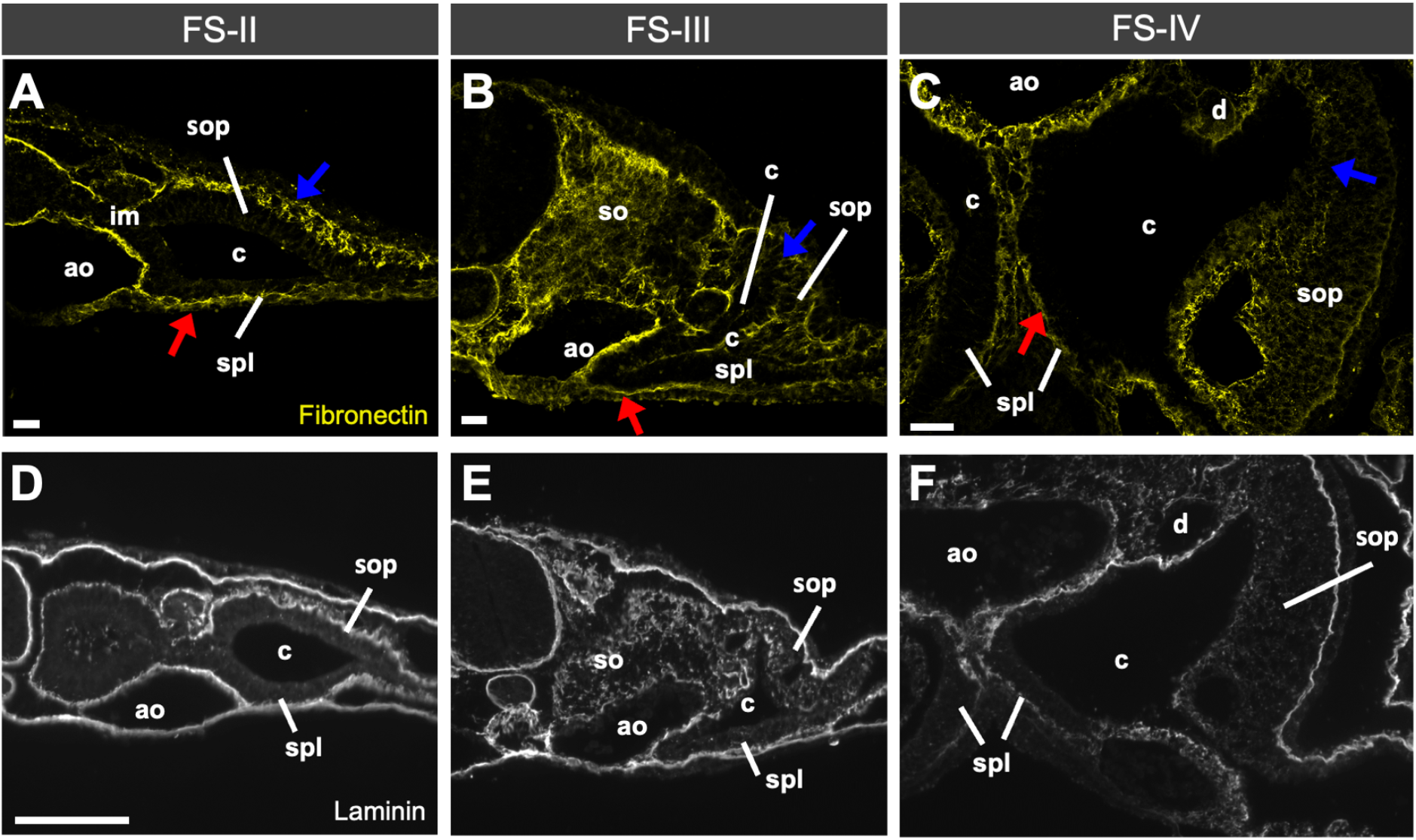
Additional changes in ECM proteins upon EMT in the Sop (supplemental to Figure 6). Sections from the indicated stages stained for Fibrillin (A-C) and Laminin (D-F). Note fragmentation of the ECM in the Sop, in contrast to maintenance of fibril morphology in the Spl.

**Supplemental Figure S2:**
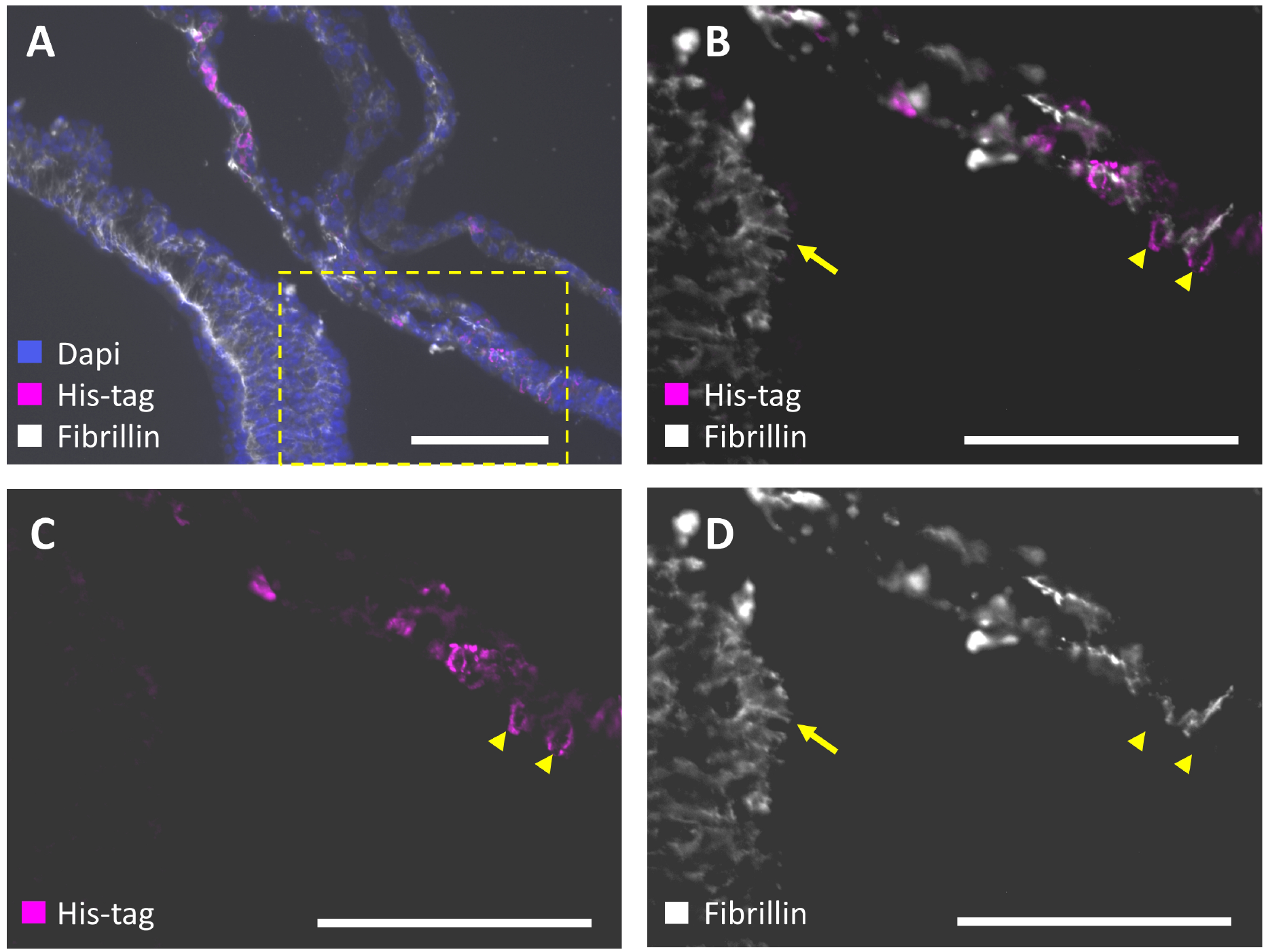
Antibody to Fibrillin-2 is chick specific (supplemental to Figure 7). Embryo was electroporated with His-tagged human Fbn2 and stained with antibodies to Fbn2 and His. The Fbn2 antibody does not colocalize with the His staining.

Movie M1: Time lapse movie of the embryo shown in Fig. 3, showing elastic recoil of the Sop and Spl after cutting the lateral connection between these layers.

